# Sea-level and climate changes drive lineage diversification in the imperiled Venus flytrap (*Dionaea muscipula* J. Ellis, Droseraceae)

**DOI:** 10.64898/2026.06.01.729438

**Authors:** Wenbin Zhou, Derick B. Poindexter, Jamie Winshell, Diego Urquia, Parul Johri, Corbin D. Jones, Gregory P. Copenhaver, Michael Kunz, John L. Randall

## Abstract

Venus flytrap (*Dionaea muscipula*) is an insectivorous plant in the monotypic genus *Dionaea*, endemic to the Southeastern United States, and found in a small geographic range in North and South Carolina. Its unique morphology and natural history have intrigued biologists for centuries, yet the evolution of populations has not been adequately characterized. Population decline, driven primarily by disruptive land conversion, fire suppression, and illegal harvesting/poaching underscores the urgency of defining the genetic diversity in the present population to guide species conservation. We applied a genome-wide SNP dataset to analyze admixture and population structure and used coalescent modeling to trace the ancestry and migration of geographically distinct population clusters. Our population dynamics model supports four lineages, with the North Cape Fear Arch lineage as the ancestor of all lineages and two genetically distinct Sandhills lineages derived from the South Cape Fear Arch lineage independently at ∼13 Ma and ∼3 Ma, respectively, during peak climate optima periods and maximal sea level rise. Severe population decline events occurred among all lineages, with two ancient instant bottleneck population decline events in Sandhills lineages and two recent ones in the two Cape Fear Arch lineages, which contributed to the current genetic structure. Shorter peduncle lengths in Sandhills populations correspond with suspected local adaptation to Sandhills habitats.

## 1 Introduction

Not until 2015 was the North American Coastal Plain recognized by Conservation International as a global biodiversity hotspot, based on the criteria that there are over 15,000 endemic taxa and over 70% habitat loss (Myers et al., 2000; Noss et al., 2015). The combined pairing of a high level of endemism with habitat loss should give a sense of urgency for understanding the evolutionary history of these taxa and prioritizing conservation actions, particularly those of acute conservation concern.

Among the most iconic taxa of the North American Coastal Plain Global Diversity Hotspot is the narrow endemic and insectivorous plant, Venus flytrap (*Dionaea muscipula* J. Ellis, Droseraceae), hereafter referred to as VFT. VFT is perhaps one of the most widely recognized plants on Earth outside of major food crops, but it is not well known that VFT is restricted to an historical 161 km (100 mi) inland radius of Wilmington, NC, USA (Coker, 1928; Hamon et al., 2021; Weakley, 2026) and is vulnerable to extinction (IUCN 2025; NatureServe 2021).

VFT occurs almost exclusively within the Cape Fear Arch (CFA) geologic formation and adjacent Sandhills physiographic province of the Carolinas, which it shares with at least 44 other endemic or near-endemic plant taxa (Leblond, 2001; Sorrie & Weakley, 2001; Ungberg et al., 2024). The ancient CFA geologic uplift occurred in the late Cretaceous period (100.5-66 Ma) and extends in a southeastward direction from near Cape Lookout, NC, to Cape Romain, SC, and northwestward beyond Fayetteville, NC, to the Sandhills physiographic province (Leblond, 2001; Marple & Hurd, 2021; Sorrie & Weakley, 2001; Walker & Coleman, 1987). The rich biological diversity and endemism contained within the CFA is largely due to areas of persistence through sea level rise events during interglacial periods (Colquhoun et al., 1991), as well as distinctive topographic complexity, oligotrophic Coastal Plain soils, and climate stability (Noss et al., 2015; Ungberg et al., 2024). Leblond (2001) provides a more thorough description of CFA geology and endemism and points out that the CFA appears to have higher native plant and animal diversity than any other area of similar size along the Atlantic Coast north of Florida. The region is also influenced by a 1-3-year fire return interval based on lightning strikes (6-10 flashes/sq. km/yr.) and purported historical anthropogenic “prescribed” fire (Frost, 2006).

A range-wide VFT Element Occurrence (EO) map for NC shows general population location, size, disposition, and overall species decline (Figure 1). Rank criteria for individual numbers per EO are defined as: A (2,000+), B (1,000-2,000), C (500-1,000), D (<500), E (documented to be present, but no specific information on number of individuals), F (failed to find), H (historical), and X (extirpated). Population decline continues (Figure S1 from North Carolina heritage program survey, 2025), largely unabated, by habitat conversion, fire suppression (Frost, 1998; Luken, 2005; Schulze et al., 2001), altered hydrology (Carter, 1975), wild collection (Evans & Gibson, 2012; Margulies et al., 2024; Sutter et al., 1982), and unintended disruption by rights-of-way (mis)management.

**Figure 1.**
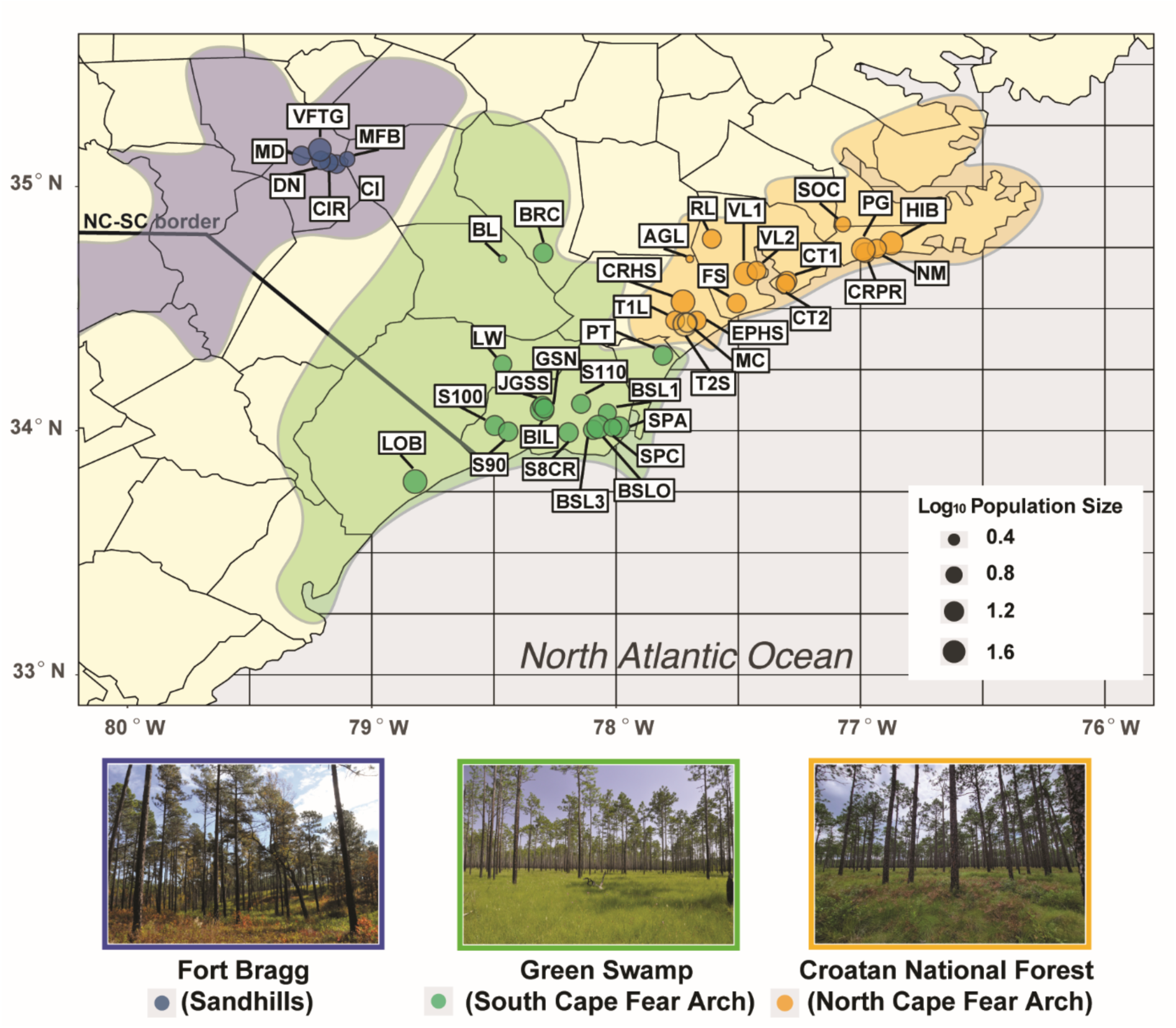
Venus flytrap (*D. muscipula*) distributions in the Carolinas, USA, across three different geographic regions: Sandhills (e.g., Fort Bragg) highlighted by blue, South Cape Fear Arch (e.g. Green Swamp) highlighted by green, and North Cape Fear Arch (e.g. Croatan National Forest) highlighted by orange. DNA from a total of 43 populations (624 accessions) were collected, with circle sizes reflecting the log 10 transformed numbers of individuals collected per population.

Phylogeographic studies that elucidate genetic structure can inform and aid conservation actions by uncovering processes that shape species’ geographic distributions, determine how migrations may have occurred over geological and generational time, serve to identify ancestral population centers, show if populations expanded or contracted and if genetic bottlenecks arose, and locate where refugial sites occurred during unfavorable times (Médail & Baumel, 2018; Schaal et al., 1998). In terms of conservation, current distributions of a species can be modeled to predict most suitable habitats in the past and for the future (Maguire et al., 2015; Peterson et al., 2011). Despite the considerable interest in VFT, which includes a detailed draft assembly of the VFT genome (Hackl, 2016; Palfalvi et al., 2020), its genetic structure and phylogeography remains undocumented.

Here, we used a large genome-wide SNP dataset generated from RAD-seq to comprehensively assess geographic structure of Venus flytrap and specifically investigated four aspects of the evolution and Venus flytrap natural history. 1) We assessed genetic diversity to reveal population structure and identify the key populations important for conservation. 2) We inferred the evolutionary history of these populations using phylogenetic and coalescence methods and explore historical and ecological factors shaping evolutionary and contemporary patterns. 3) We used species distribution modeling (SDM) to predict the most suitable habitats for VFT (especially for the Sandhills populations) and inform conservation strategies.

## 2 Materials and Methods

### 2.1 Study Species

Taxonomically, VFT is in a monotypic genus with no close relatives in North America (Weakley, 2025). It is not found growing naturally nor does it have a pollen record outside of its restricted area of the Carolinas (Delcourt & Delcourt, 1981; Sorrie & Weakley, 2001). Thus, although capable of colonizing other suitable habitats (where planted in the USA states of Florida, Delaware, and New Jersey), VFT only naturally inhabits the current and restricted region within which it evolved. Moreover, VFT occurs in low pH soils and has a very distinctive tetrahedral tetrad pollen morphology (Halbritter et al., 2012) which would most likely allow for ready identification if present in pollen cores.

*Dionaea* spp. microfossil pollen does occur, however, in the middle Miocene (23-5.3Ma) and Pliocene (5.3-2.6 Ma) of central Europe that is comparable to modern *Dionaea muscipula* (Muller, 1981) indicating that this genus, or perhaps an extinct relative, was once more common and widespread in distribution than today (Cameron et al., 2002). VFT’s closest relative, *Aldrovanda vesiculosa* (Waterwheel Plant) is an Old-World taxon (Europe, Africa, Southeast Asia, and Australia), strictly aquatic, with a submerged snap-trap prey-capturing mechanism, and was similarly more widespread than today (Kundu et al., 1996). Darwin (1875) even remarked that *Aldrovanda* was ‘‘a miniature, aquatic *Dionaea*.’’

On the natural community level, VFT habitat includes Wet Pine Flatwoods and Savannas, Streamhead Pocosins, Canebrakes, Small Depression Shrub Borders, Carolina Bays, ecotones between pocosins and more upland Longleaf Pine/Scrub Oak savannas, and Sandhills Seeps (NatureServe, 2021; Schafale, 2024; Weakley, 2026). VFT also occurs in unintentional refugia associated with human-created ditches, fire line breaks, roadsides, and other managed rights-of-way (Hamon et al. 2021; US Fish and Wildlife Service 2023).

### 2.2 Sample collection and DNA preparation

We collected 624 VFT tissue samples from 43 populations across its native range in the Carolinas, USA (Figure 1; Table S1). These accessions covered three different geographic regions, including the Sandhills, the South Cape Fear Arch (SCFA), and North Cape Fear Arch (NCFA) (Figure 1). We separated the two areas for geographic comparison based on division north and south of the Cape Fear River. Populations targeted for tissue collection were selected based on physical distance (> 2km) to maximize the ability to detect genetic variation across the range. Three to five leaves were collected using sanitized scissors from each plant and stored on dry ice or at -80°C before DNA extraction.

DNA was isolated using the Plant DNAzol kit (Chomczynski et al., 1997). DNA quality was assessed using UV light visualization of ethidium bromide staining after electrophoresis on 1% agarose gel and quantified using a NanoDrop spectrophotometer (Thermo Fisher Scientific, USA) and Quant-it PicoGreen dsDNA kit (Invitrogen, USA).

In addition to VFT tissue collection, a companion project was conducted to collect, process, and store VFT seeds as a population genome library. Over 94K seeds, from 65 sites, were collected by maternal line, frozen at -20^0^ C in vacuum-sealed foil laminate bags and stored in the North Carolina Botanical Garden seed bank. These seeds are to ensure against extinction of wild populations not under conservation protection, and to provide a resource for research and legitimate reintroduction/augmentation projects.

### 2.3 RAD-Seq

A total of 200 ng DNA of each sample was digested with EcoRI-HF (New England Biolabs, Ipswich, MA, USA; NEB) and *Nla*III (NEB) in Cutsmart buffer (NEB). DNA fragments were barcoded by Illumina indexes and barcodes (see details in Table S1). In total, seven libraries were prepared. DNA samples from the same library were pooled proportionally, followed by the size-selection step using Agilent 4200 TapeStation (Agilent, Santa Clara, CA, USA) to capture DNA fragments ranging from 250 to 950 bp. Each pooled library was amplified for 12 PCR cycles using the Illumina indexed primer. Subsequently, the pooled libraries were loaded to HiSeq 4000 (Illumina, Inc., San Diego, CA, USA) for 50 bp paired ends sequencing, respectively. Library preparation and sequencing were carried out at the High Throughput Sequencing Facility at University of North Carolina at Chapel Hill.

### 2.4 Raw Data processing

Samples were demultiplexed first using customized bash scripts for the different barcodes and indexes listed in Table S1. Then, all sorted samples were trimmed, clustered, filtered, and aligned using ipyrad (Eaton & Overcast, 2020). Due to lack of high-quality genome assembly, we generated our loci clusters based on *de novo* assembly following Zhou et al. (2018). In the filtering step, we kept loci shared at least in 230 samples (over 53% final kept samples), which could generate the relatively best phylogeny with relatively high total number of loci and more good quality samples according to (Zhou et al., 2018; Zhou & Xiang, 2022). Missing data, caused by allele dropout, can influence downstream phylogeographic analyses (Gautier et al., 2013), so we only kept samples with less than 40% missing loci in their final RAD-seq data matrix (431 individuals in total, Table S1). All analyses were done on the Longleaf cluster at UNC-Chapel Hill.

### 2.5 Genetic Diversity and Population Structure

To assess patterns of genetic diversity and differentiation among populations, we carried out nucleotide diversity (π) estimation, Analysis of Molecular Variance analysis (AMOVA) (Excoffier et al., 1992), principal component analysis (PCA) (Abdi & Williams, 2010), and STRUCTURE (Pritchard et al., 2000) analysis based on the final RAD-seq data matrix.

We calculated π for each population. Due to the small sample sizes after the filtering step, populations with less than three individuals were combined with the population geographically nearest to them. Specifically, we merged one BL individual into the BRC population and one AGL individual into the RL population. Pi was estimated for each population and each geographic region to evaluate within and among-population genetic variation based on all RAD-seq loci shared in over 90% individuals. The analysis was performed using vcftools (Danecek et al., 2011) to get the full alignments and a customized Python script to estimate the π using the traditional equation –

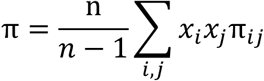

(Nei, 1987; Nei & Li, 1979; Tajima, 1983). We did not consider the degenerated sequences as a polymorphism in the python script. We then assessed genetic variation among geographic regions and populations using (AMOVA) in poppr from the R package (Kamvar et al., 2014). This hierarchical analysis tested the proportion of genetic variance attributed to among-region, among-population, and within-population levels. We used 999 permutations for AMOVA based on the filtered SNP matrix.

We performed PCA on the RAD-seq data to visualize genetic variation among individuals and to assess if clustering patterns correspond to geographic patterns. PCA was conducted following the tutorial from ipyrad analysis tool using the filtered SNP matrix as an input and population names (Table S1). We then plotted the 3D PCA using the plotly function in R (Sievert, 2019) with top three principal components as x, y, and z axis, respectively.

Finally, we applied a model-based clustering approach, structure, using ezstructure, which is a repackaged version of fastSTRUCTURE (Raj et al., 2014) to estimate individual ancestry proportions and infer the number of genetically distinct clusters (K) for VFT. We ran ezstructure in fastSTRUCTURE with default priors and convergence settings across a range of K values from 2 to 10. After model fitting, we used the structure_choosek function in ezstructure to determine the optimal K value that maximizes marginal likelihood. We then visualized the best K value structure plot using ezdistruct function in ezstructure.

### 2.6 Phylogenetic analysis

To detect the evolutionary patterns among lineages of VFT, we conducted phylogenetic analyses using IQ-TREE v2 (Minh et al., 2020) based on the final filtered RAD-seq dataset (M50 matrix). This M50 data matrix includes all loci present in at least 50% of the 431 individuals and excludes samples with more than 40% missing data (including Ns and gaps) to ensure high data quality. Maximum likelihood trees were inferred using “GTR+G” model. Branch support was evaluated using 1,000 ultrafast bootstrap replicates. While *Aldrovanda* and *Dionaea* are both carnivorous snap-trap plants within the family Droseraceae, they diverged approximately 45 million years ago (Palfalvi et al., 2020). Such long time means that substantial genetic differences have accumulated in *Aldrovanda,* which cannot serve as a reliable root for analyzing the recent population-level history of VFT. Therefore, the tree was rooted from the crown node of monophyletic North Cape Fear Arch populations. The rooting strategy was later tested using fastsimcoal2 (Excoffier et al., 2021). The resulting consensus tree was visualized using iTOL (Letunic & Bork, 2021), with node support values greater than 95 displayed on each branch and all population labels highlighted at the tips. To confirm the relationships among populations, we also used SplitsTree4 (Huson & Bryant, 2024) to construct a split network from the M50 unlinked SNP dataset, employing the NeighborNet algorithm with variance calculated by ordinary least squares.

### 2.7 Isolation by Distance Test

To test whether an island model or a stepping-stone model (Kimura, 1953) might explain the current distribution of VFT, we used an isolation-by-distance method (Wright, 1943). We first calculated pairwise F_ST_ values (Meirmans & Hedrick, 2011) as a measure of genetic distance between populations, based on the SNP dataset using Weir & Cockerham method (Weir & Cockerham, 1984) and geographic distances (km) based on their coordinates of populations using the geosphere package in R (Hijmans et al., 2017). Subsequently, a Mantel test was performed to evaluate the correlation between the genetic distance matrix (F_ST_ / (1 – F_ST_)) and the geographic distance matrix, using 9999 permutations to assess statistical significance using the vegan package in R (Dixon, 2003). The results were drawn by ggplot2 in R (Wickham, 2016). Because Sandhills populations were isolated from the SCFA and NCFA populations, we also investigated the correlation between genetic distance and geographic distance within each geographic region.

### 2.8 Sea Level Changes along the US East Coast

From the literature, we assembled patterns of sea level changes on the eastern coast of the United States, including North Carolina and neighboring states. The geologic literature is clear that there was a major shift of sea level on the eastern coast of North Carolina (Rowley et al., 2013). We also extracted climate data and timing from (Miller et al., 2020; Johnson et al., 2021). These data were then compared to our simulation results (section 2.9).

### 2.9 Population Dynamics

To infer historical population dynamics, including changes of population size and lineage-sorting time, we applied fastsimcoal2 v2.8 (Excoffier et al., 2021), a coalescent-based simulation framework based on fitting the site frequency spectrum (SFS). We constructed a multi-dimensional folded joint SFS from the filtered SNP data which includes RAD-seq loci shared in at least 90% of individuals using *easySFS.py* (https://github.com/isaacovercast/easySFS#easysfs) for three and four lineages according to geographic regions (Figure 1) and genetic structures (Figure 2), respectively.

**Figure 2.**
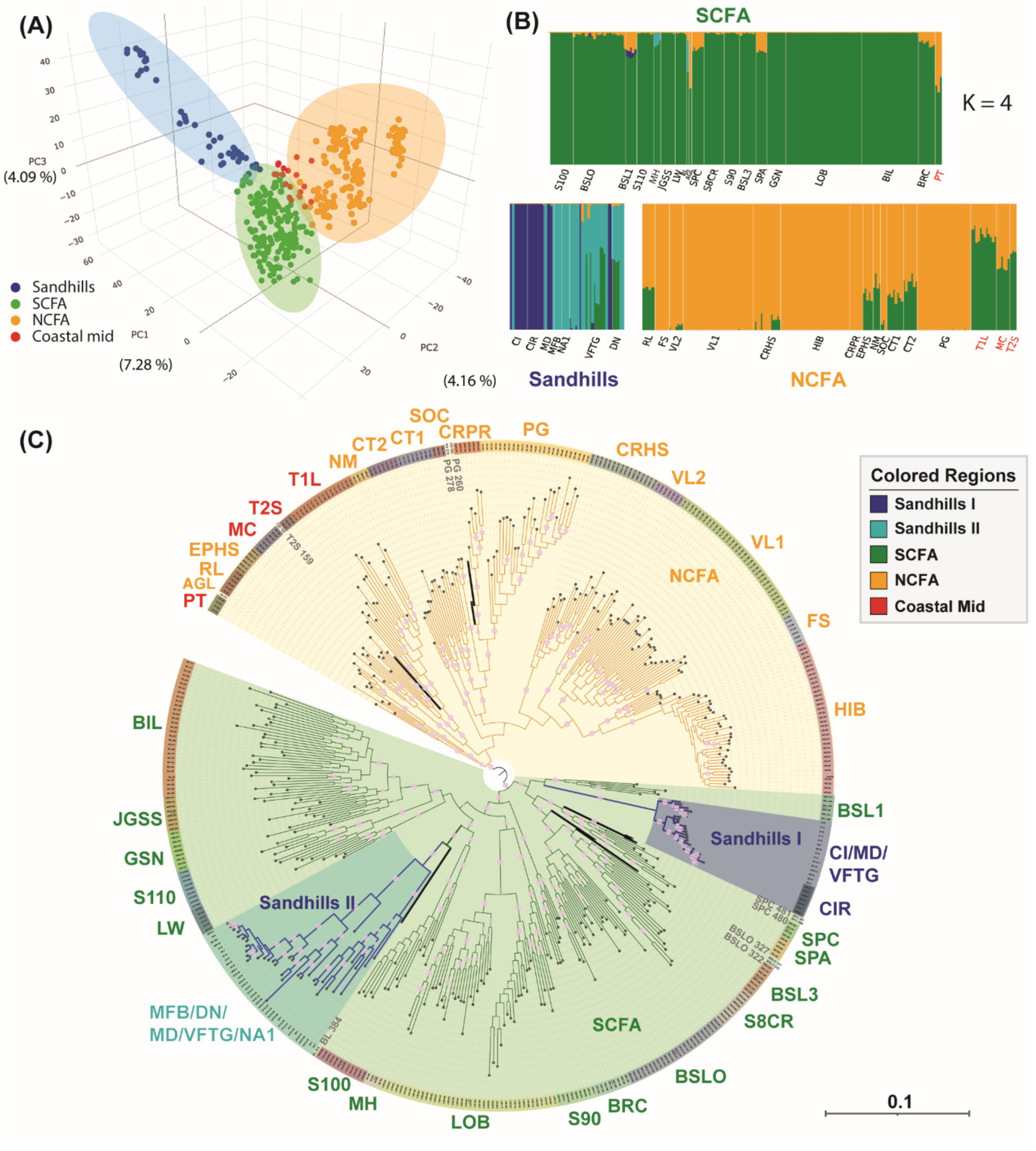
Population structure and phylogeny of Venus flytrap (VFT) indicate current populations were derived from four ancestral lineages and match well with their geographic distributions. (A) 3D Principal Component Analysis (PCA) of VFT based on from RAD-seq genetic data indicates three major clusters matching to geographic locations, including South Cape Fear Arch (SCFA) (Green), North Cape Fear Arch (NCFA) (Orange), and Sandhills (Blue) in the Carolinas. The red dots indicate four populations located at boundary of SCFA and NCFA, which are also genetically clustered between SCFA and NCFA clusters. It includes PT, T2S, T1L, and MC populations. (B) Structure analysis of VFT (best K = 4) suggests four major ancestral gene pools. SCFA Populations shared one ancestral gene pool (green color), NCFA populations derived from one ancestral gene pool (orange color), and Sandhills populations consist of two different ancestral gene pools (light and dark blue color). Populations in red font match with the coastal middle populations, indicating a highly mixed gene pool. (C) The phylogeny of VFT detects a complicated evolutionary history. NCFA populations form a monophyletic clade, which is sister to the clade of SCFA populations and Sandhills populations. Notably, the Sandhills populations consist of two monophyletic clades derived from SCFA populations at different times. Most populations in NCFA and SCFA are monophyletic, which are highlighted by different colors at tips. However, all populations in Sandhills are paraphyletic except CIR population. Individuals in gray color and thickened lines are paraphyletic, including BSLO, SPC, T2S, and PG. All pink dots at each node present the UFBS values greater than 95.

We started with the simplest models to test whether each lineage underwent historical bottleneck events. We specifically tested each lineage with four scenarios, including no effective population size (*Ne*) change, with a single instantaneous bottleneck (estimating the current population size, bottleneck intensity, and time of bottleneck), and a single size change both instantly and exponentially (estimating the current population size and growth/decline rate in both cases). The model with an instant size change was found to best fit the data for all geographic populations and lineages (Figure S2), with a decline inferred in all lineages. All parameters are provided in Github (https://github.com/Bean061/venus_flytrap_phylogeography/tree/main/5fastsimcoal2/github/final_run_1pop/) and the estimated parameters are provided in Table S2.

Corresponding to the three geographic distributions (Figure 1) – Sandhills, South Cape Fear Arch (SCFA), and North Cape Fear Arch (NCFA), we fitted a three-population demographic model. Each population was assumed to undergo an instantaneous single-size change model (as referred to be the best model when using single population as described above). Due to the lack of a closely related outgroup for phylogenetic analysis, the ancestral population is unknown. Therefore, we (1) considered the SCFA lineage as the ancestral lineage, followed by the formation of Sandhills lineages, and the formation of NCFA lineage independently (model “3Pop_M1”); (2) assumed the NCFA lineage as the ancestor, followed by the SCFA lineage and Sandhills lineages independently (model “3Pop_M2”); and (3) regarded the Sandhills lineages as the ancestor, followed by the SCFA and NCFA lineages independently (model “3Pop_M3”) (Figure S3). The first two models lead to similar low AIC values (Figure S3D) while the third model had a much higher AIC value and was thus excluded. A total of 22 parameters were estimated in these three-population models (Table S3).

Next, corresponding to the STRUCTURE results, four-population models were fitted to the lineages NCFA, SCFA, and two lineages from the Sandhills. Again, based on the best-fit single-population models, each population was assumed to undergo an instantaneous single-size change model. Based on the top two models from the three-lineage scenarios, we considered the models with NCFA or SCAF as the ancestor. More specifically, the following two models were fitted: (4) NCFA lineage was set as the ancestor, followed by formation of SCFA lineage and two Sandhills lineages independently (model “4Pop_M1”); and (5) SCFA populations were set as the ancestral lineage, followed by formation of two Sandhills lineages and the NCFA lineage independently (model “4Pop_M2”) (Figure S4). A total of 26 parameters were estimated (Table S3). The best fit model supports NCFA lineage as the ancestor (“4Pop_M1”) with lowest AIC value compared to the other four-population models (Figure S4C) and all three-population models (Figure S3D).

We selected the best-fit model from the four-lineage scenarios (“4Pop_M1”) and modified the model with 10 different gene flow scenarios across different time periods (see a schematic of the models considered in Figure S5). A total of 27 - 34 parameters estimated in these four-population models (Table S4). We set the mutation rate at 6.95 x 10^-9^ per site per generation in the same way as the well-studied model species, *Arabidopsis thaliana* (L.) Heynh. (Weng et al., 2019). For each model, *fastsimcoal2* was run with 15 independent replicates, each using 100,000 coalescent simulations, and parameter optimization conducted with the Expectation-Conditional Maximization (ECM) algorithm over 40 loops. Each model was repeated 15 times. The best-fit model was selected based on AIC values. All relevant scripts for all models are available in Github (https://github.com/Bean061/venus_flytrap_phylogeography/tree/main/5fastsimcoal2/).

### 2.10 Niche Comparison and Species Niche Modeling

To evaluate the niche variance of VFT among the three geographic regions, we employed PCA analysis using 20 climatic variables at 2.5-minutes resolution from WorldClim v2.0 (Fick & Hijmans, 2017), and 15 soil variables at 2.5 arc seconds from SoilGrids 2.0 (Poggio et al., 2021). To stack all layers at the same resolution and scale, we applied the resample and crop function in the R terra package. We then included occurrence records from the Global Biodiversity Information Facility (GBIF, www.gbif.org) and our own field collections. From the GBIF dataset we retained only unique, wild-occurring records that clearly fell within the species’ native range to reduce the influence of cultivated or misidentified samples. To reduce the analysis biases from unbalanced numbers of occurrences from each geographic region, we also cleaned occurrences by keeping one occurrence within a 5km geographic distance circle (2.5 arc minutes).

To reduce interactions among colinear variables and obtain a more accurate prediction of VFT distribution, we used a variable inflation factor method to remove multicollinear variables before environmental niche modeling (ENM) following (Rautsaw et al., 2022). After filtering, a total of 18 variables were kept for ENM, including eight WorldClim variables and ten SoilGrids variables. We then conducted SDM analysis using the MaxEnt algorithm with both linear and quadratic models implemented in the R ENMevaluate package (Muscarella et al., 2014). The model with the highest value area under the receiver operating characteristic curve (AUC) was plotted (DeLong et al., 1988). To identify the most suitable current VFT habitat we conducted ENM using all known VFT occurrences from the Carolinas. Due to the particularly xeric environment in Sandhills, we also tested whether the Sandhills populations might lead to a different environmental niche using Sandhills occurrences only. We lastly conducted PCA analyses and an ANOVA test to verify the niche differentiation among three geographic regions using VIF filtered environmental variables.

### 2.11 Potential Fitness Trait Comparison

To test for local adaptation among populations, we compared a potential fitness-related trait—peduncle length—among VFT specimens. Ninety-eight digitized herbarium specimens from the University of North Carolina Herbarium (NCU) via SERNEC (SouthEast Regional Network of Expertise and Collections) were each assigned to one of three geographic regions based on county of origin. Peduncle length was measured using ImageJ (Abràmoff et al., 2004), and differences among the three regions were evaluated using an ANOVA test.

## 3 Results

### 3.1 RAD-seq captures genetic variation across populations

We generated approximately 2,300 million raw reads for 624 individuals with an average of ∼3.8 million reads per individual. After filtering individuals with more than 40% missing data, we kept a total of 431 individuals with an average of ∼4.8 million reads for downstream analyses. The final dataset covered all 43 populations with 32,588 loci (24,809 loci per accession) and 146,908 SNPs (Table S1).

### 3.2 Genetic diversity and population structure analyses reveal four main VFT lineages across three geographic regions

VFT nucleotide diversity (π) indicated an overall low but similar diversity across all 43 populations with a mean of 0.001 (Table S5). Among three geographic regions, π values in NCFA, SCFA, and Sandhills are 0.0016, 0.0018, and 0.0020, respectively. Interestingly, the Sandhills had the lowest population sizes based on survey data but possessed two populations with especially high π values (e.g., 0.0018 in population CI and 0.0020 in population DN (Table S6). The AMOVA analysis based on SNP data shows a significant differentiation (*P* < 0.001) among lineages or geographic VFT regions with a high percentage of variance explained among populations (52.48% when K = 4 and 39.94% when K = 3 in Table 1).

**Table 1.**
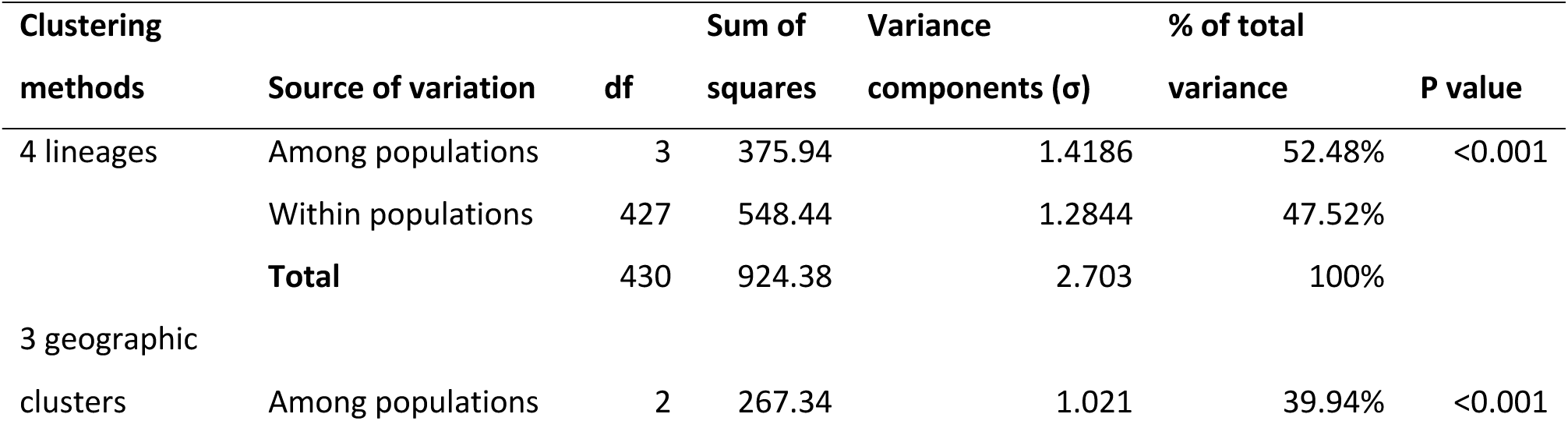
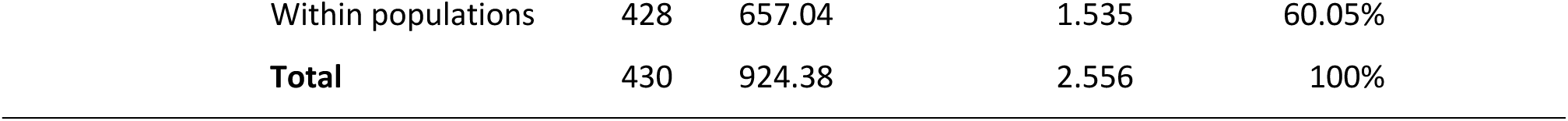
AMOVA test results from 999 permutations among different population clusters.

Mean Fixation Index (Fst) between paired populations is 0.315 (Table S6), and the highest F_ST_ value (0.758) was observed between CIR (Sandhills) and SOC (NCFA) (Table S5). The top two genetically distinguished populations were CIR from the Sandhills and HIB from the most Northeast population in the NCFA with an average F_ST_ value of 0.520 and 0.494 with other populations (Table S6). Interestingly, the previously mentioned three most diverged populations (CIR, SOC, and HIB) had the lowest π values, with 0.0002, 0.0004, and 0.0004, respectively (Table S5).

The 3D-PCA results based on RAD-seq SNP data indicated populations from the three regions were differentiated by the top three principal components. Specifically, Sandhills populations (both light and dark blue color dots) were differentiated from all Coastal populations (green and orange dots in Figure 2A). Within the coastal populations, a putative hybridization zone occurred between SCFA and NCFA, and share a similar genetic background based on PCA analysis. These admixed populations include PT, T2S, T1L, and MC, (red dots in Figure 2A). Some admixed populations exist, however, without overlapping contact (i.e., BSL1, SPC, SPA in SCFA; EPHS, NM, CT1, CT2, and RL in NCFA; and VFTG and DN in Sandhills). The overall PCA results mirrored the structure analysis (Figure 2A, 2B & Figure S6), suggesting populations from the same geographic regions largely share the same major ancestral genetic components. The best-fit model supported K equals 4 (Figure 2B) and revealed two major ancestral gene pools in the Sandhills region rather than the expected one.

Phylogenetic analyses using IQ-TREE and SplitTree4, based on the M50 dataset of 431 VFT accessions, revealed that the Sandhills populations are nested within the Cape Fear Arch populations (Figure 2C & Figure S7). The IQ-TREE results indicated that NCFA populations formed a monophyletic group, while Cape Fear Arch populations were paraphyletic. The two Sandhills lineages were apparently derived from the Cape Fear Arch lineage and independently formed two monophyletic clades (Figure 2C), where the Sandhills I is sister to the BSL population and Sandhills II is sister to the BL population. The SplitTree4 analysis supported the same monophyly pattern of the NCFA populations and the paraphyly of the Cape Fear Arch populations. Although all Sandhills lineages appeared to cluster together, the CIR, MD, and CI populations from the Sandhills II lineage were more closely related to the PT and BRC populations in the Cape Fear Arch populations (Figure S7).

### 3.3 Sea level change helped shape population history of VFT

The isolation by distance (IBD) analysis using all 43 populations found significant but modest correlation between genetic and geographic distances among all populations, partially supporting the “stepping-stone” theory (Mantel test, r = 0.5104, P < 0.001, Figure 3). This consistent pattern suggests that limited gene flow tends to occur among geographically close populations. However, populations with similar genetic distance between 0.25 and 0.6 (Figure 3A) occur across a wide geographic range (> 200 km). All within IBD correlation disappears within each geographic region (All P > 0.05). This indicates that, besides the stepping-stone model, other historical dispersal drivers or bottleneck events have likely contributed to the current VFT distribution.

**Figure 3.**
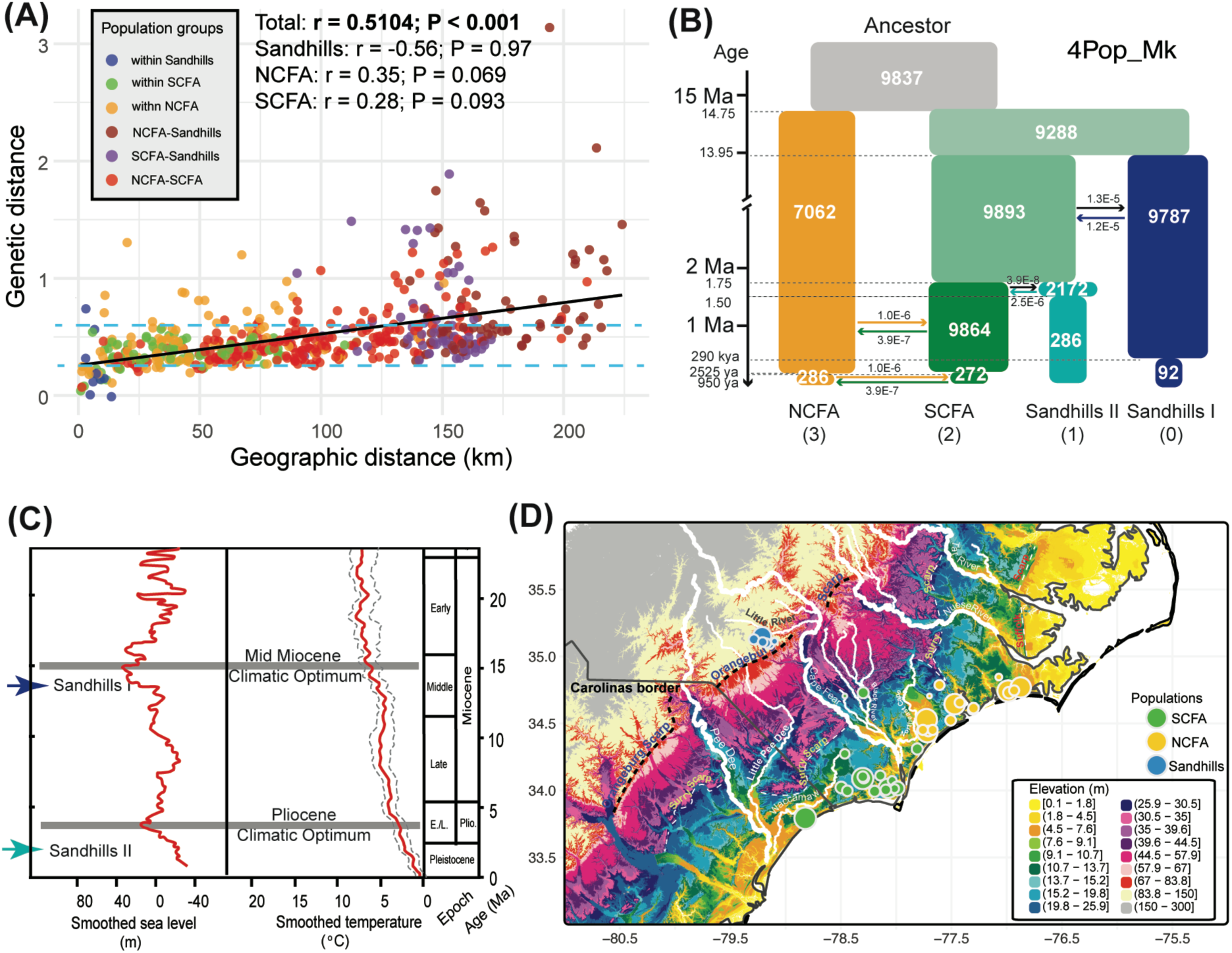
Population dynamics of Venus Flytrap suggest that historical climate changes and resulting sea level changes might have shaped the current distribution. A) IBD analysis reveals a significantly modest correlation between Genetic distance and Geographic distance (r = 0.5104, *P* < 0.001). However, within each geographic region, the correlations become non-significant. The two blue dashed lines indicate the populations with genetic distance between 0.25 and 0.6 can occur at a great distance. B) The best-fit model from *fastsimcoal2* indicates a formation time of current lineages traced back to 14.75 Ma. A four-lineage scenario suggests four instant population decline events occurred, with two recent ones in NCFA and SCFA lineages at ∼2500 years ago, while two ancient ones in Sandhills lineages at ∼1.5 Ma and ∼0.29 Ma, respectively. It also detected recent gene flow between NCFA and SCFA as well as ancient gene flow between Sandhills and SCFA lineages. Numbers in each box represent population size (x1000). C) Two formation time of Sandhills lineages are around the two Climatic Optima. Left box indicates projected sea level change and the right box indicate the projected temperature since Miocene by Johnson (2021) and Miller et al. (2020). D) Lidar image of eastern Coast of Carolinas indicates the sea level changes. The Orangeburg scarp indicates the Sandhills regions were historically at the coastal line. SH represents Sandhills, SCFA represents South Cape Fear Arch, and NCFA represents North Cape Fear Arch.

Assuming a five-year VFT generation time (based on the approximate flowering time from our greenhouse materials), the best-fit four-lineage model (Model “4Pop_Mk”, Figure 3B) from *fastsimcoal2* indicated that a VFT ancestor split into the North Cape Fear (NCFA) lineage and the SCape Fear Arch (SCFA) lineage approximately 15 million years ago (Ma), or 2.95 million generations ago (mga) (Figure S5F). The two Sandhills lineages were then derived from the SCFA lineage at 13.95 Ma (2.79 mga) and 1.75 Ma (0.35 mga), respectively (Figure 3B). The dates for the formation time of the Sandhills lineages match well with dates derived from geological data for the two climatic optima and sea level changes since the Miocene (Figure 3C, D).

Our analyses also detected instant population size decline in each lineage (Figure 3B), and our one-lineage scenario supported instant population decline size over bottleneck events and expansional size change events (Figure S2). Two instant population size decline events occurred at approximately 1.5 Ma and 0.29 Ma in Sandhills I and Sandhills II, respectively, while two recent ones occurred in coastal lineages approximately 2500 years ago (Figure 3B). In addition, our hypothetical models presume different gene flow events among populations (Figure S5) and suggest that gene flow occurred between the Sandhills lineages and SCFA lineages before population decline events within Sandhills lineages (Figure 3B) presumably when Sandhills lineages were isolated and gene flow was restricted. Gene flow between SCFA and NCFA lineages is more recent and persisted since ∼1.75 Ma (Figure 3B).

### 3.5 Sandhills populations appear differentiated in niche and morphology

Although environmental niche model (ENM) could be limited when climatic data is not fully represent, geographic ranges are restricted, and population sizes are small (Qiao et al., 2017; Saupe et al., 2012), the optimal ENM, including all occurrences of VFT, predicted the most suitable habitats are along the Carolinas’ coast with extremely low suitability in the Sandhills region (Figure 4A). Suitable habitat prediction changed dramatically and was directed inland when only using input from Sandhills occurrences (Figure 4B). Moderate niche differentiation is inferred between the Sandhills and coastal populations. PCA analysis, based on VIF variables, confirms that Sandhills populations are substantially different from coastal populations (Figure 4C). Major variables differentiating the Sandhills and coastal populations include one soil variable (Coarse Fragments), three precipitation related variables (Bio13 Precipitation of Wettest Month, Bio14 Precipitation of Driest Month, and Bio19 Precipitation of Coldest Quarter), two temperature related variables (Bio7 Temperature Annual Range and Bio9 Mean Temperature of Driest Quarter), and Elevation (Figure 4C).

**Figure 4.**
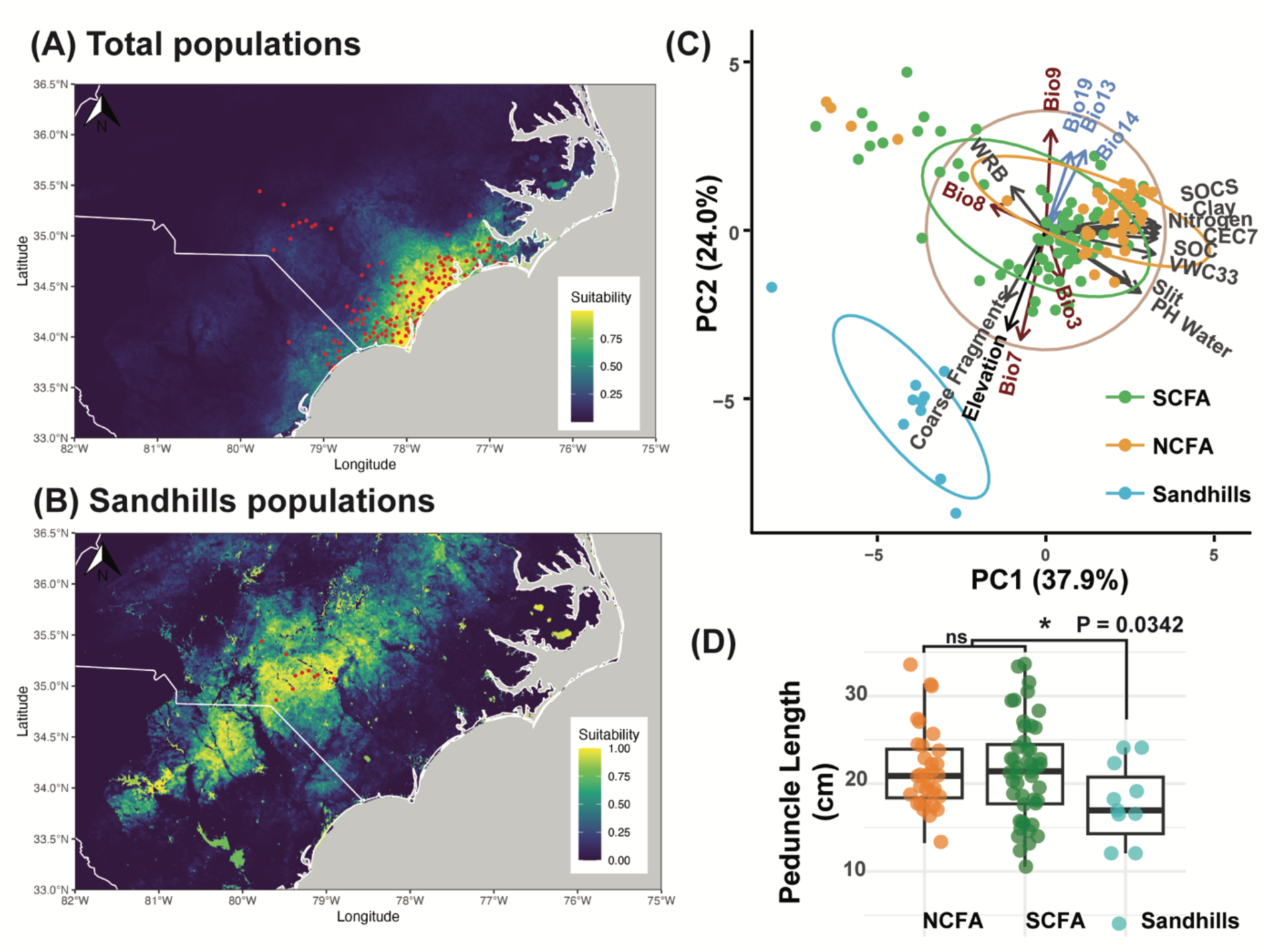
Differentiation of Niche and Morphology indicates a certain level of local adaptation in VFT. (A) ENM using all current occurrences indicates the most suitable habitats are on the coastal regions of the Carolinas. (B) ENM using the current Sandhills occurrences indicates the most suitable habitats of Sandhills lineages are inland of the Carolinas. (C) The PCA analysis indicates niche differentiation between Sandhills and two coastal lineages. Variables in red are related to temperature, including Bio3 Isothermality, Bio7 Temperature Annual Range, Bio8 Mean Temperature of Wettest Quarter, and Bio9 Mean Temperature of Driest Quarter. Variables in blue are related to precipitation, including Bio13 Precipitation of Wettest Month, Bio14 Precipitation of Driest Month, and Bio19 Precipitation of Coldest Quarter. All gray variables are related to soils, including WRB (World Reference Base), SOC (Soil organic carbon), SOCS (Soil organic carbon stock), CEC7 (Cation exchange capacity at PH 7), VWC33 (volume of water content at -33 kPa), Clay, Nitrogen, Silt, Water PH. (D) Peduncle lengths in Sandhills lineages are significantly lower than the ones in coastal lineages. Different letters indicate the significant ad hoc test following ANOVA. NCFA represents North Cape Fear Arch, SCFA represents South Cape Fear Arch, and SH represents Sandhills.

Similarly, morphological analyses of peduncle length revealed phenotypic differences between Sandhills and coastal populations, consistent with patterns of population isolation. Individuals from both coastal populations have significantly higher peduncle lengths than Sandhills populations (ANOVA, P = 0.0342) based on herbariums specimens (Figure 4D).

When integrating genetic, climatic, and morphological data, a key pattern is concordant: genetically distinct populations occupy divergent climatic niches and possess different physiological traits. The alignment between population genetic structure, environmental gradients, and trait variation suggests that local adaptation has shaped genetic divergence (in at least one trait) among populations after populations became isolated due to sea level changes.

## 4 Discussion

### 4.1 Venus flytrap phylogeographic patterns as driven by ancient sea level changes

Our data shows that the extant distribution of VFT and the genetic differentiation among these populations is in part the legacy of past changes in sea-level in the Carolinas. Beginning with the Early Eocene (∼53 Ma), sea level of the U.S. Atlantic Coastal Plain changed dramatically across the glacial maxima and climate optimal periods (Cronin et al., 1981; Krantz, 1991). Along the U.S. Atlantic Coastal Plain, particularly in North Carolina, sea-level rise was (and is) much faster than the eastern coast of the Atlantic Ocean (Kemp et al., 2011; Long et al., 2014). This accelerated rise reshaped low-elevation habitats, shorelines, and foredune morphologies (Jay et al., 2022). These changes were driven by shifted groundwater tables, altered soil moisture gradients, and changes in disturbance regimes that affected the biotic community (reviewed by Powell et al., 2017).

Coastal population of VFT (NCFA and SCFA) are both tied to the CFA, an ancient Mid Eocene (45 Ma) geological formation and an active area of continuing uplift (Van De Plassche et al., 2014). Portions of the CFA likely remained exposed across repeated periods of inundation, presumably providing a terrestrial refugium for VFT and many other taxa now endemic to this region (Colquhoun et al. 1991; LeBlond 2001). From the Mid Miocene (∼15 Ma) to the present, VFT distribution expanded and contracted at least twice where the outer coastal plain of the Carolinas varied from 80–120 km west of the current coastline to the full width of the continental shelf. The Last Glacial Maximum (∼18 Ka) shoreline would have been located at the shelf edge, adding approximately 3K sq km of the potential VFT habitat to the coastal plain (Thieler et al., 2014). Additionally, during the past 10,000 years, VFT might have had a much more extensive distribution and a more complicated phylogeography and genetic diversity than we presently see.

### 4.2 Population structure not fully explained by isolation by distance

In our data we identified four major ancestral lineages of VFT (Figure 2). These correspond to three broad geographic regions, with two distinct lineages that emerged from coastal lineages (SCFA and NCFA) and two that emerged from the Sandhills region. Population dynamics modeling and phylogeny suggest that these lineages originated at different times. The ancestral populations diverged into the NCFA and SCFA lineages about 15 Ma, while the two Sandhills lineages appear to have diverged independently from the SCFA ancestral lineage at ∼1.75 Ma and ∼14.75 Ma, respectively (Figures 2C and 3B). These divergence times, which were derived solely from the population genetic data, align strikingly well with two climatic optima - the Pliocene climatic optimum around 3 Ma and the Middle Miocene climatic optimum around 15 Ma (Figure 3C; Johnson, 2021; Miller et al., 2020) when sea levels reached their peaks and submerged large portions of current coastal regions (Dumitru et al., 2019; Rowley et al., 2013) other than exposed refugial areas of the CFA.

Coastal populations from SCFA and NCFA regions share overlapping primarily habitats, but independent ranges (Figure 2C). As noted, the Sandhills also have two distinct populations. Our isolation by distance result shows a significant correlation between the genetic distance and geographic distance across all populations, but the coefficient value is moderate (r= 0.51; Fig. 3). Therefore, geographic isolation does explain some of the patterns of population differentiation in our data. However, as shown in Figure 3A, within each geographic region the coefficient values are lower and not statistically significant, suggesting that within these populations VFT genetic differences are not explained by the isolation by distance model. However, as shown in Table 1 (and Table S6), there remains considerable genetic differentiation within each of these populations. This suggests that other factors may drive VFT genetic differentiation within these populations.

### 4.3 Formation of greater Cape Fear Arch region populations

The NCFA/SCFA population cluster regions are likely differentiated by the Northeast Cape Fear River barrier and associated floodways, river terraces, and other riparian habitats. The NCFA cluster, formed approximately 14 Ma, represents an ancestral lineage and supports the idea that the greater CFA axial high remained emergent through the Mid Miocene and Pliocene climate optima, and during subsequent sea-level rise (van de Plassche et al., 2014). Of course, the habitat VFT occupied prior to the Holocene is unknown, although it likely was open with ample sunlight within areas of wet oligotrophic soils. While fire frequency was more prevalent only within the last 10,000 years (Delcourt & Delcourt, 1981), it is also reasonable to believe that herbivorous megafauna such as giant ground sloths, mastodons, bison, and horses of the future Carolinas (Sanders, 2002) could have also helped keep VFT habitat free of overtopping vegetation during the Quaternary.

Despite the distinctiveness of the NCFA and SCFA lineages, our results suggest introgression between them. Several populations contain an admixed structure with both NCFA and SCFA ancestral components. These populations include those located near the geographic boundary between the two regions (e.g., PT, T2S, MC, EPHS, CRHS, T1L, SPC, BSL1, and SPA), including Holly Shelter game land and south of Cape Fear River system, suggesting that these areas may represent an historical or ongoing secondary contact zone between the two lineages.

Interestingly, the admixture pattern is also observed in several populations in the northern NCFA region (e.g., AGL, BRC, RL, VL2, NM, CT1, CT2, and SOC), but not in many southern SCFA populations (Figure 2B). The complicated admixture pattern suggests that introgression may not be restricted to the current contact zone between SCFA and NCFA. Instead, this pattern may indicate historical gene flow that occurred when these ancestral lineages expanded following the receding of the coastline. Consistent with these observations, the demographic inferences suggest introgression between SCFA and NCFA occurred relatively recently (< 1.75 Ma). Because NCFA regions presumably remained exposed since the late Cretaceous (Allman et al., 2016; Figure S8), these glacial CFA refugia could have allowed for SCFA and NCFA lineage hybridization during sea level and climate oscillations.

### 4.4 Formation of Sandhills populations

The Sandhills VFT populations are ecologically distinct from coastal populations and occur in Streamhead Pocosins and Sandhill Seeps, where sphagnum moss tends to co-occur, and where other endemic and near-endemic plants occur in the larger Sandhills ecosystem (Sorrie, 2016). As noted above, these populations likely became isolated because of sea-level rise. Our results show that two distinct genetic lineages formed (Sandhills I and Sandhills II), both derived from SCFA, but at two different times (∼13.5 Ma and ∼1.75 Ma, respectively) and with limited gene flow (Figures. 1 and 2B). The Sandhills region formed in the Cretaceous (Marple & Hurd, 2021) but was overtopped by a younger geological formation through eolian processes ∼70 kya (Swezey et al., 2016). The divergent Sandhills habitats may also have influenced population structure and bottleneck events. Given that VFT seeds are slightly larger (1–3 mm) than coarse sand fragments (0.35-0.59 mm), strong northwest eolian winds may have facilitated dispersal of VFT seeds toward the SCFA and suitable habitat (based on stepping-stone models) could plausibly explain the mixed genetic components observed in coastal-margin populations such as MH and BL (Figure 2B).

The persistence of Sandhills populations supports the hypothesis that VFT has adapted to this distinct niche. Within the Sandhills populations each creek or stream system share the same ancestral gene pools (e.g., CI–CIR and VFTG–DN), implying long-term lineage persistence of these populations and little to no admixture between the two Sandhills lineages (Figure 2b). In this environment, isolation of these populations may be driven by unsuitable intervening habitat or barriers to pollination (Hamon et al., 2019; Youngsteadt et al., 2018). However, the unadmixed structure within the same population at distances of less than 2 km (i.e., VFTG, DN) remains unknown.

The Sandhills populations tolerate an ecologically distinct niche that is strongly differentiated by abiotic factors (Figure 4). Additionally, the VFT in the Sandhills are morphologically distinct; significantly shorter peduncle lengths occur in Sandhills populations compared to the coastal populations (Figure 4D). A recent common-garden experiment demonstrated that Sandhills populations do exhibit significantly higher growth rates than coastal populations under controlled water, nutrition, and light treatments (Morris et al., in prep.). All the above evidence suggests that morphological differences might reflect underlying genetic differentiation of these populations.

### 4.5 Smaller scale dispersal drivers may shape the current distribution of Venus Flytrap

Large-scale phenomena such as sea level rise and river systems clearly shaped the evolutionary history and geographic distribution of VFT. Other smaller scale factors could also be driving subtle structure in each VFT population. Seed dispersal of the small (1-2mm diameter) seeds is highly localized and caused by gravity and rainwater sheet flow. Long-distance dispersal is not considered an ordinary mechanism and is probably exceedingly rare. However, there is documented seed capsule herbivory by white-tailed deer and preliminary evidence that VFT seeds can survive intact through the white-tailed deer digestive tract (J. Randall, personal observation). While rare, if these long-range dispersal events did lead to the founding of new populations when the sea level receded, this could result in localized population differentiation.

### 4.6 Range-wide VFT conservation implications

The extremely low nucleotide diversity (≈0.001) among almost all VFT populations indicate that current populations are incredibly vulnerable (Table S5). Similar π values were also found in many other endangered species using genome wide SNPs, e.g. *Gingko biloba* (≈0.002) (Zhao et al., 2019), *Cupressus chengiana* (≈0.008) (Li et al., 2020). Isolation by distance across all 43 populations analyzed suggests limited gene flow even between geographically proximate populations. The northwestward extension of the SCFA to the Sandhills and Piedmont was likely once a likely migratory corridor that could have mitigated the negative effects of population isolation, such as inbreeding and stochastic population decline through attrition, but are now faced with barriers in the current fragmented landscape.

Potential gene flow between and among populations is likely pollinator driven rather than via seed dispersal. Known primary pollinator species – small sweat bees *Augochlorella gratiosa* and *Lasioglossum creberrimum*, and beetles *Typocerus sinuatus* and *Trichodes apivorus* – typically forage only locally (Youngsteadt et al., 2018; Greenleaf et al., 2007). We therefore suggest that all extant VFT populations – particularly those that occur in sites not under conservation protection or maintained by best management practices – receive special conservation attention. This is perhaps particularly urgent for Sandhills populations (with their genetic uniqueness) and the smaller and/or peripheral populations where, for example, inbreeding and genetic drift could drive them toward local extinction or reduce the potential for environmental change response.

We also believe that VFT, as both a Flagship and Umbrella species, can directly raise awareness for both conservation efforts and protection of the larger ecological community within which it occurs. Many other endemic and near-endemic plant taxa also warrant phylogeographic studies in the Cape Fear Arch proper and Sandhills. VFT and its associated taxa are also part of a global ecologically related plant alliance of “Old, Climatically Buffered, Infertile Landscapes” – OCBILs – reviewed by Hopper (2009) and allied with other centers of vascular plant endemism.

Conservation of these ancient and ecologically allied taxa, like VFT, that likely found refuge in either the exposed areas of the Cape Fear Arch during sea level rise episodes or in the Sandhills through northwestern arm and migration corridor of the SCFA (LeBlond 2001; Sorrie 2013) is essential to preserving the genetic legacy of these remarkably resilient species.

## ACKNOWLEDGEMENTS

This work was supported by the International Carnivorous Plant Society with the award number A16-1891-001. Special thanks to Carol Ann McCormick and Shannon Oberreiter for kindly providing the digitized images of all Venus flytrap vouchers from the NCU herbarium. Thanks to Bruce Sorrie and Eric Ungberg for discussions on the plant species of the Sandhills, to Alan S. Weakley for his insight into VFT phylogeography and regional plant endemism, Richard J. LeBlond for an early review of the manuscript, and to Todd J. Vision, Allen Hurlbert, Feng Xiao, Jeffrey Thorne, Xiang Ji, and Fay-Wei Li for Methods discussions. Particular thanks go to the organizations, agencies, and land managers that provided collecting permits and the many individuals who carefully collected VFT tissue (and seeds in a related project) across the species’ range. Tyler Gramley is especially acknowledged for providing the index plant on which we practiced DNA extractions, sequencing, and assembly.

## AUTHOR CONTRIBUTIONS

John L. Randall is the PI of this project; he initiated the idea, obtained funding, coordinated the collection of materials, and contributed to writing. Wenbin Zhou conducted all data analyses with Derick Poindexter and helped draft the first version of manuscript. Jamie Winshell organized the DNA extraction and sequencing. Diego Urquia did pilot data analyses and proof the final draft. Parul Johri provided important guidance on fastsimcoal2 methods and proofread the population genomics sections. Corbin D. Jones and Gregory P. Copenhaver co-authored the International Carnivorous Plant Society funding proposal and multiple manuscript reviews. Michael Kunz assisted with coordinating VFT seed and tissue collection and manuscript review.

## DATA AVAIL ABILITY STATEMENT

All multiplexed ddRADseq data have been deposited in the NCBI Sequence Read Archive (SRA) under BioProject ID PRJNA1276505 with the accession numbers SAMN49074409–SAMN49074460. All scripts are available on Github: https://github.com/Bean061/venus_flytrap_phylogeography.git.

## REFERENCES

1. Abdi, H., & Williams, L. J. (2010). Principal component analysis. Wiley Interdisciplinary Reviews: Computational Statistics, 2(4), 433–459.

2. Abràmoff, M. D., Magalhães, P. J., & Ram, S. J. (2004). Image processing with ImageJ. Biophotonics International, 11(7), 36–42.

3. Cameron, K. M., Wurdack, K. J., & Jobson, R. W. (2002). Molecular evidence for the common origin of snap-traps among carnivorous plants. American Journal of Botany, 89(9), 1503–1509. 10.3732/ajb.89.9.1503

4. Carter, L. J. (1975). Agriculture: A new frontier in coastal North Carolina. Science, 189(4199), 271–275.

5. Chomczynski, P., Mackey, K., Drews, R., & Wilfinger, W. (1997). DNAzol®: A reagent for the rapid isolation of genomic DNA. Biotechniques, 22(3), 550–553.

6. Coker, W. C. (1928). THE DISTRIBUTION OF VENUS’S FLY TRAP (DIONAEA MUSCIPULA). Journal of the Elisha Mitchell Scientific Society, 43(3–4), 221–228.

7. Colquhoun, D.J., G.H. Johnson, P.C. Peebles, P.F. Huddleston, and T. Scott, 1991: Quaternary geology of the Atlantic Coastal Plain. In: Quaternary Nonglacial Geology: Conterminous

8. U.S. [Morrison, R.B. (ed.)]. The Geology of North America v. K-2. Geological Society of America, Boulder, CO, pp. 629–650.

9. Cronin, T. M., Szabo, B. J., Ager, T. A., Hazel, J. E., & Owens, J. P. (1981). Quaternary Climates and Sea Levels of the U.S. Atlantic Coastal Plain. Science, 211(4479), 233–240. 10.1126/science.211.4479.233

10. Danecek, P., Auton, A., Abecasis, G., Albers, C. A., Banks, E., DePristo, M. A., Handsaker, R. E., Lunter, G., Marth, G. T., Sherry, S. T., & others. (2011). The variant call format and VCFtools. Bioinformatics, 27(15), 2156–2158.

11. Darwin, C. (1875). Insectivorous plants. J. Murray.

12. Delcourt, P. A., & Delcourt, H. R. (1981). Vegetation maps for eastern North America: 40,000 yr BP to the present. Geobotany II, 123–165.

13. DeLong, E. R., DeLong, D. M., & Clarke-Pearson, D. L. (1988). Comparing the Areas under Two or More Correlated Receiver Operating Characteristic Curves: A Nonparametric Approach. Biometrics, 44(3), 837–845. JSTOR. 10.2307/2531595

14. Dixon, P. (2003). VEGAN, a package of R functions for community ecology. Journal of Vegetation Science, 14(6), 927–930.

15. Dumitru, O. A., Austermann, J., Polyak, V. J., Fornós, J. J., Asmerom, Y., Ginés, J., Ginés, A., & Onac, B. P. (2019). Constraints on global mean sea level during Pliocene warmth. Nature, 574(7777), 233–236.

16. Eaton, D. A., & Overcast, I. (2020). ipyrad: Interactive assembly and analysis of RADseq datasets. Bioinformatics, 36(8), 2592–2594.

17. Evans R., T. Gibson, and Y.B. Johnson. 2012. Conservation Status of Venus Flytraps. In Tim Bailey and Stewart McPherson, editors. Dionaea: The Venus’s Flytrap. Dorset (England): Redfern Natural History Productions. p. 127–141.

18. Excoffier, L., Marchi, N., Marques, D. A., Matthey-Doret, R., Gouy, A., & Sousa, V. C. (2021). fastsimcoal2: Demographic inference under complex evolutionary scenarios. Bioinformatics, 37(24), 4882–4885.

19. Excoffier, L., Smouse, P. E., & Quattro, J. M. (1992). Analysis of molecular variance inferred from metric distances among DNA haplotypes: Application to human mitochondrial DNA restriction data. Genetics, 131(2), 479–491.

20. Fick, S. E., & Hijmans, R. J. (2017). WorldClim 2: New 1-km spatial resolution climate surfaces for global land areas. International Journal of Climatology, 37(12), 4302–4315. 10.1002/joc.5086

21. Frost, C. (2006). History and future of the longleaf pine ecosystem. *The Longleaf Pine Ecosystem: Ecology*, Silviculture, and Restoration, 438, 9–42.

22. Frost, C. C. (1998). Presettlement fire frequency regimes of the United States: A first approximation. Fire in Ecosystem Management: Shifting the Paradigm from Suppression to Prescription. Tall Timbers Fire Ecology Conference Proceedings, 20, 70–81.

23. Gautier, M., Gharbi, K., Cezard, T., Foucaud, J., Kerdelhué, C., Pudlo, P., Cornuet, J.-M., & Estoup, A. (2013). The effect of RAD allele dropout on the estimation of genetic variation within and between populations. Molecular Ecology, 22(11), 3165–3178.

24. Greenleaf, S. S., Williams, N. M., Winfree R., and Kremen C. (2007). Bee foraging ranges and their relationship to body size. Oecologia (2007) 153:589–596.

25. Hackl, T. (2016). A draft genome for the Venus Flytrap, Dionaea muscipula: Evaluation of assembly strategies for a complex Genome–Development of novel approaches and bioinformatics solutions [PhD Thesis]. Dissertation, Würzburg, Universität Würzburg, 2016.

26. Halbritter, H., Hesse, M., & Weber, M. (2012). The unique design of pollen tetrads in Dionaea and Drosera. Grana, 51(2), 148–157.

27. Hamon, L. E., Youngsteadt, E., Irwin, R. E., & Sorenson, C. E. (2019). Pollination Ecology and Morphology of Venus Flytrap in Sites of Varying Time Since Last Fire. Annals of the Entomological Society of America, 112(3), 141–149. 10.1093/aesa/say032

28. Hamon, L., Hannon, D., Mason, S., Horton, S., & Buchanan, M. (2021). A RANGEWIDE STATUS SURVEY OF VENUS FLYTRAP DIONAEA MUSCIPULA(DROSERACEAE).

29. Hijmans, R. J., Williams, E., Vennes, C., & Hijmans, M. R. J. (2017). Package ‘geosphere.’ Spherical Trigonometry, 1(7), 1–45.

30. Hopper, S. D. (2009). OCBIL theory: Towards an integrated understanding of the evolution, ecology and conservation of biodiversity on old, climatically buffered, infertile landscapes. Plant and Soil, 322(1), 49–86.

31. Huson, D. H., & Bryant, D. (2024). The SplitsTree App: Interactive analysis and visualization using phylogenetic trees and networks. Nature Methods, 21(10), 1773–1774.

32. IUCN. (2025). The IUCN Red List of Threatened Species. Version 2025-2. https://www.iucnredlist.org. Accessed on [2 March 2025].

33. Jay, K. R., Hacker, S. D., Hovenga, P. A., Moore, L. J., & Ruggiero, P. (2022). Sand supply and dune grass species density affect foredune shape along the US Central Atlantic Coast. Ecosphere, 13(10), e4256. 10.1002/ecs2.4256

34. Johnson, M. E. (2021). Geological oceanography of the Pliocene warm period: A review with predictions on the future of global warming. Journal of Marine Science and Engineering, 9(11), 1210.

35. Kamvar, Z. N., Tabima, J. F., & Grünwald, N. J. (2014). Poppr: An R package for genetic analysis of populations with clonal, partially clonal, and/or sexual reproduction. PeerJ, 2, e281.

36. Kemp, A. C., Horton, B. P., Donnelly, J. P., Mann, M. E., Vermeer, M., & Rahmstorf, S. (2011). Climate related sea-level variations over the past two millennia. Proceedings of the National Academy of Sciences, 108(27), 11017–11022. 10.1073/pnas.1015619108

37. Kimura, M. (1953). “ Stepping-stone” model of population.

38. Krantz, D. E. (1991). A chronology of Pliocene sea-level fluctuations: The US Middle Atlantic Coastal Plain record. Quaternary Science Reviews, 10(2–3), 163–174.

39. Kundu, S., Basu, S., & Chakraverty, R. (1996). Aldrovanda vesiculosa Linn.-It’s maiden appearance and disappearance from India. Journal of Economic and Taxonomic Botany, 20, 719–724.

40. Leblond, R. J. (2001). Endemic Plants of the Cape Fear Arch Region. Castanea, 66(1–2), 83–97.

41. Letunic, I., & Bork, P. (2021). Interactive Tree Of Life (iTOL) v5: An online tool for phylogenetic tree display and annotation. Nucleic Acids Research, 49(W1), W293–W296.

42. Li, J., Milne, R.I., Ru, D., Miao, J., Tao, W., Zhang, L., Xu, J., Liu, J., Mao, K. (2020). Allopatric divergence and hybridization within *Cupressus chengiana* (Cupressaceae), a threatened conifer in the northern Hengduan Mountains of western China. Molecular Ecology, 29(7), 1250–1266. 10.1111/mec.15407

43. Long, A. J., Barlow, N. L. M., Gehrels, W. R., Saher, M. H., Woodworth, P. L., Scaife, R. G., Brain, M. J., & Cahill, N. (2014). Contrasting records of sea-level change in the eastern and western North Atlantic during the last 300 years. Earth and Planetary Science Letters, 388, 110–122. 10.1016/j.epsl.2013.11.012

44. Luken, J. O. (2005). Habitats of Dionaea muscipula (Venus’ Fly Trap), Droseraceae, Associated with Carolina Bays. Southeastern Naturalist, 4(4), 573–584. 10.1656/1528-7092(2005)004%255B0573:HODMVF%255D2.0.CO;2

45. Maguire, K.C., Nieto-Lugilde, D., Fitzpatrick, M.C., Williams, J.W., Blois, J.L. (2015). Modeling species and community responses to past, present, and future episodes of climatic and ecological change. Annual Review of Ecology, Evolution, and Systematics, 46(1), 343–368. 10.1146/annurev-ecolsys-112414-054441

46. Margulies, J. D., Trost, B., Hamon, L., Kerr, N. Z., Kunz, M., Randall, J. L., Shew, R. D., Shew, D. M., Starke, L., Suiter, D., & others. (2024). Expert assessment of illegal collecting impacts on Venus flytraps and priorities for research on illegal trade. Conservation Biology, 38(5), e14320. 10.1111/cobi.14320

47. Marple, R. T., & Hurd, Jr., J. D. (2021). Investigation of the Cape Fear arch and East Coast fault system in the Coastal Plain of North Carolina and northeastern South Carolina, USA, using LiDAR data. Atlantic Geology, 57, 311–341. 10.4138/atlgeol.2021.015

48. Médail, F., & Baumel, A. (2018). Using phylogeography to define conservation priorities: The case of narrow endemic plants in the Mediterranean Basin hotspot. Biological Conservation, 224, 258–266. 10.1016/j.biocon.2018.05.028

49. Meirmans, P. G., & Hedrick, P. W. (2011). Assessing population structure: *F*_ST_ and related measures. Molecular Ecology Resources, 11(1), 5–18. 10.1111/j.1755-0998.2010.02927.x

50. Miller, K. G., Browning, J. V., Schmelz, W. J., Kopp, R. E., Mountain, G. S., & Wright, J. D. (2020). Cenozoic sea-level and cryospheric evolution from deep-sea geochemical and continental margin records. Science Advances, 6(20), eaaz1346.

51. Minh, B. Q., Schmidt, H. A., Chernomor, O., Schrempf, D., Woodhams, M. D., Von Haeseler, A., & Lanfear, R. (2020). IQ-TREE 2: New models and efficient methods for phylogenetic inference in the genomic era. Molecular Biology and Evolution, 37(5), 1530–1534.

52. Muller, J. (1981). Fossil pollen records of extant angiosperms. The Botanical Review, 47(1), 1.

53. Muscarella, R., Galante, P. J., Soley-Guardia, M., Boria, R. A., Kass, J. M., Uriarte, M., & Anderson, R. P. (2014). ENMeval: An R package for conducting spatially independent evaluations and estimating optimal model complexity for Maxent ecological niche models. Methods in Ecology and Evolution, 5(11), 1198–1205. 10.1111/2041-210X.12261

54. Myers, N., Mittermeier, R. A., Mittermeier, C. G., Da Fonseca, G. A. B., & Kent, J. (2000). Biodiversity hotspots for conservation priorities. Nature, 403(6772), 853–858. 10.1038/35002501

55. NatureServe. (2021). NatureServe Explorer [web application]. NatureServe, Arlington, Virginia. Available https://explorer.natureserve.org/Taxon/ELEMENT_GLOBAL.2.159781/Dionaea_muscipula. (Accessed: March 2, 2021).

56. Nei, M. (1987). Molecular evolutionary genetics. Columbia university press.

57. Nei, M., & Li, W.-H. (1979). Mathematical model for studying genetic variation in terms of restriction endonucleases. Proceedings of the National Academy of Sciences, 76(10), 5269–5273.

58. Noss, R. F., Platt, W. J., Sorrie, B. A., Weakley, A. S., Means, D. B., Costanza, J., & Peet, R. K. (2015). How global biodiversity hotspots may go unrecognized: Lessons from the North American Coastal Plain. Diversity and Distributions, 21(2), 236–244. 10.1111/ddi.12278

59. Palfalvi, G., Hackl, T., Terhoeven, N., Shibata, T. F., Nishiyama, T., Ankenbrand, M., Becker, D., Förster, F., Freund, M., Iosip, A., Kreuzer, I., Saul, F., Kamida, C., Fukushima, K., Shigenobu, S., Tamada, Y., Adamec, L., Hoshi, Y., Ueda, K., … Hedrich, R. (2020). Genomes of the Venus Flytrap and Close Relatives Unveil the Roots of Plant Carnivory. Current Biology, 30(12), 2312–2320.e5. 10.1016/j.cub.2020.04.051

60. Peterson, A.T., Soberón, J., Pearson, R.G., Anderson, R.P., Martínez-Meyer, E., Nakamura, M., Araújo, M.B., (2011). Ecological niches and geographic distributions. In Ecological niches and geographic distributions. Princeton university press.

61. Poggio, L., de Sousa, L. M., Batjes, N. H., Heuvelink, G. B. M., Kempen, B., Ribeiro, E., & Rossiter, D. (2021). SoilGrids 2.0: Producing soil information for the globe with quantified spatial uncertainty. SOIL, 7(1), 217–240. 10.5194/soil-7-217-2021

62. Powell, E. J., Tyrrell, M. C., Milliken, A., Tirpak, J. M., & Staudinger, M. D. (2017). A synthesis of thresholds for focal species along the U.S. Atlantic and Gulf Coasts: A review of research and applications. Ocean & Coastal Management, 148, 75–88. 10.1016/j.ocecoaman.2017.07.012

63. Pritchard, J. K., Stephens, M., & Donnelly, P. (2000). Inference of population structure using multilocus genotype data. Genetics, 155(2), 945–959.

64. Qiao, H., Peterson, A.T., Ji, L., Hu, J. (2017). Using data from related species to overcome spatial sampling bias and associated limitations in ecological niche modelling. Methods in Ecology and Evolution, 8(12), 1804–1812. 10.1111/2041-210X.12832

65. Raj, A., Stephens, M., & Pritchard, J. K. (2014). fastSTRUCTURE: variational inference of population structure in large SNP data sets. Genetics, 197(2), 573–589.

66. Rautsaw, R. M., Jiménez-Velázquez, G., Hofmann, E. P., Alencar, L. R. V., Grünwald, C. I., Martins, M., Carrasco, P., Doan, T. M., & Parkinson, C. L. (2022). VenomMaps: Updated species distribution maps and models for New World pitvipers (Viperidae: Crotalinae). Scientific Data, 9(1), 232. 10.1038/s41597-022-01323-4

67. Rowley, D. B., Forte, A. M., Moucha, R., Mitrovica, J. X., Simmons, N. A., & Grand, S. P. (2013). Dynamic topography change of the eastern United States since 3 million years ago. Science, 340(6140), 1560–1563.

68. Sanders, A. E. (2002). Additions to the Pleistocene mammal faunas of south Carolina, north Carolina, and Georgia. American Philosophical Society.

69. Saupe, E.E., Barve, V., Myers, C.E., Soberón, J., Barve, N., Hensz, C.M., Peterson, A.T., Owens, H.L., Lira-Noriega, A. (2012). Variation in niche and distribution model performance: the need for a priori assessment of key causal factors. Ecological Modelling, 237, 11–22. 10.1016/j.ecolmodel.2012.04.001

70. Schaal, B. A., Hayworth, D. A., Olsen, K. M., Rauscher, J. T., & Smith, W. A. (1998). Phylogeographic studies in plants: Problems and prospects. Molecular Ecology, 7(4), 465–474. 10.1046/j.1365-294x.1998.00318.x

71. Schafale, M. P. (2024). Classification of the Natural Communities of North Carolina Fourth Approximation. North Carolina Natural Heritage Program N.C. Department of Natural and Cultural Resources Raleigh, NC

72. Schulze, W., Schulze, E., Schulze, I., & Oren, R. (2001). Quantification of insect nitrogen utilization by the venus fly trap Dionaea muscipula catching prey with highly variable isotope signatures. Journal of Experimental Botany, 52(358), 1041–1049.

73. Sievert, C. (2019). *Plotly for R* [Computer software]. https://plotly-r.com.

74. Sorrie, B. A. (2016). The Curious Distribution of Asclepias tomentosa (Apocynaceae). Phyton, 68, 1–4.

75. Sorrie, B. A., & Weakley, A. S. (2001). Coastal Plain Vascular Plant Endemics: Phytogeographic Patterns. Castanea, 66(1–2), 50–82.

76. Sutter, R. D., Mowbray, T. B., & May, M. L. (1982). The status and collection of Venus Flytrap (Dionaea muscipula) in North Carolina. North Carolina Plant Conservation Program, The North Carolina Department of Agriculture, Plant Industry Division.

77. Swezey, C. S., Fitzwater, B. A., Whittecar, G. R., Mahan, S. A., Garrity, C. P., González, W. B. A., & Dobbs, K. M. (2016). The Carolina Sandhills: Quaternary eolian sand sheets and dunes along the updip margin of the Atlantic Coastal Plain province, southeastern United States. Quaternary Research, 86(3), 271–286.

78. Tajima, F. (1983). Evolutionary relationship of DNA sequences in finite populations. Genetics, 105(2), 437–460.

79. Thieler, E. R., Foster, D. S., Himmelstoss, E. A., & Mallinson, D. J. (2014). Geologic framework of the northern North Carolina, USA inner continental shelf and its influence on coastal evolution. Marine Geology, 348, 113–130. 10.1016/j.margeo.2013.11.011

80. Ungberg, E. A., Horn, J. W., Poindexter, D. B., Bradley, K. A., & Weakley, A. S. (2024). Helianthus waccamawensis (Asteraceae), a new species of sunflower endemic to the Cape Fear Arch Region of North and South Carolina (U.S.A.). Journal of the Botanical Research Institute of Texas, 18(2). 10.17348/jbrit.v18.i2.1365

81. U.S. Fish and Wildlife Service. (2023). Species status assessment report for the Venus Flytrap (*Dionaea muscipula*). Version 1.1. March 2023. Atlanta, GA

82. Van De Plassche, O., Wright, A. J., Horton, B. P., Engelhart, S. E., Kemp, A. C., Mallinson, D., & Kopp, R. E. (2014). Estimating tectonic uplift of the Cape Fear Arch (south-eastern United States) using reconstructions of Holocene relative sea level. Journal of Quaternary Science, 29(8), 749–759. 10.1002/jqs.2746

83. Walker, H. J., & Coleman, J. M. (1987). Atlantic and Gulf coastal province.

84. Ward, H. T., & Davis, R. S. (1999). Time before history: The archaeology of North Carolina. UNC Press Books.

85. Weakley, A.S., and Southeastern Flora Team. (2026). Flora of the southeastern United States Web App. University of North Carolina Herbarium, North Carolina Botanical Garden, Chapel Hill, U.S.A. https://fsus.ncbg.unc.edu/main.php?pg=show-taxon-detail.php&lsid=urn:lsid:ncbg.unc.edu:taxon:{0E7150DB-AFF6-4D05-AE79-C440311B5745}. Accessed May 5, 2026.

86. Weir, B. S., & Cockerham, C. C. (1984). Estimating F-statistics for the analysis of population structure. Evolution, 1358–1370.

87. Weng, M.-L., Becker, C., Hildebrandt, J., Neumann, M., Rutter, M. T., Shaw, R. G., Weigel, D., & Fenster, C. B. (2019). Fine-Grained Analysis of Spontaneous Mutation Spectrum and Frequency in Arabidopsis thaliana. Genetics, 211(2), 703–714. 10.1534/genetics.118.301721

88. Wickham, H. (2016). Getting Started with ggplot2. In Ggplot2: Elegant graphics for data analysis (pp. 11–31). Springer.

89. Wright, S. (1943). Isolation by distance. Genetics, 28(2), 114.

90. Youngsteadt, E., Irwin, R. E., Fowler, A., Bertone, M. A., Giacomini, S. J., Kunz, M., Suiter, D., & Sorenson, C. E. (2018). Venus Flytrap Rarely Traps Its Pollinators. The American Naturalist, 191(4), 539–546. 10.1086/696124

91. Zhao, Y.P., Fan, G., Yin, P.P., Sun, S., Li, N., Hong, X., Hu, G., Zhang, H., Zhang, F.M., Han, J.D., Hao, Y.J. (2019). Resequencing 545 ginkgo genomes across the world reveals the evolutionary history of the living fossil. Nature Communications, 10(1), 4201. 10.1038/s41467-019-12133-5

92. Zhou, W., Ji, X., Obata, S., Pais, A., Dong, Y., Peet, R., & Xiang, Q.-Y. J. (2018). Resolving relationships and phylogeographic history of the Nyssa sylvatica complex using data from RAD-seq and species distribution modeling. Molecular Phylogenetics and Evolution, 126, 1–16.

93. Zhou, W., & Xiang, Q.-Y. J. (2022). Phylogenomics AND biogeography of Castanea (chestnut) and Hamamelis (witch-hazel)–Choosing between RAD-seq and Hyb-Seq approaches. Molecular Phylogenetics and Evolution, 176, 107592.

